# Refinement of locomotor activity is developmentally regulated by dopaminergic signaling in larval zebrafish

**DOI:** 10.1101/2025.10.27.684440

**Authors:** Briee Mercier, Sandra M. Garraway, Matthew L. Beckman, Mark A. Masino

## Abstract

The refinement of gross motor skills, such as locomotion, during development is conserved across vertebrate species. Our previous work demonstrated, in larval zebrafish, that dopaminergic signaling through the dopamine D2-like family of receptors, specifically the dopamine 4 receptor subtype, was necessary for the developmental transformation of behaviorally relevant locomotor activity from an immature to a mature pattern between 3- and 4-days post-fertilization. In this study, we used a complement of tools, including electrophysiology, pharmacology, *in vivo* calcium imaging, liquid chromatography-mass spectrometry, and quantitative reverse transcription polymerase chain reaction to characterize the functional and molecular mechanisms responsible for this dopaminergic-mediated refinement of spinal locomotor activity. The results demonstrate that the dopamine 4 receptor subtype is both present and functional in, at least, a subset of immature larvae. Further, gene expression of all D2-like receptor subtypes, levels of dopamine, and activity of diencephalic dopaminergic neurons are significantly greater in mature larvae compared to immature larvae. The integration of these results provides evidence for the developmental role of dopaminergic signaling, specifically the dopamine receptor 4 subtype, in the refinement of locomotor activity in vertebrates.

**Significance Statement:** Throughout life, all vertebrates acquire and improve gross motor skills. This is particularly evident in the locomotor system where motor output is initially coarse and becomes progressively more refined during development. Previously, we demonstrated that dopaminergic signaling was a factor in the developmental refinement of locomotor activity. However, an understanding of the molecular and functional mechanisms underlying the dopaminergic-mediated refinement of spinal locomotor activity remains elusive. This study demonstrates, in larval zebrafish, that increased expression of all D2-like dopamine receptor subtypes, levels of dopamine, and activity of diencephalic dopaminergic neurons correlate with the refinement of locomotor activity.

## Introduction

Throughout life all vertebrates acquire and improve gross motor skills. This is particularly evident in the locomotor system where motor output is initially coarse and becomes progressively more refined during development. For example, a developmental transformation that refines locomotor activity in larval zebrafish occurs between 3- and 4-days post-fertilization (dpf) (Buss and Drapeau, 2001; Brustein et al., 2003; Lambert et al., 2012). Swimming in larval zebrafish is composed of discrete episodes separated by periods of inactivity (Buss and Drapeau, 2001). At an immature stage of development (3 dpf), larval zebrafish are mostly inactive, and swimming is composed of erratic, long duration (∼1000 msec) episodes. In contrast, at a more mature stage of development (≥ 4 dpf), larval zebrafish are more active, and swimming is composed of goal-directed, short duration (∼250 msec) episodes. Our previous work demonstrated that dopaminergic signaling through inhibitory D2-like receptor (D2-likeR) subtypes (D2R, D3R, D4R), specifically D4R, was necessary for the developmental transformation of this behaviorally relevant locomotor activity (Lambert et al., 2012). However, an understanding of the molecular and functional mechanisms underlying the dopaminergic-mediated refinement of spinal locomotor activity remains elusive.

In vertebrates, the dopaminergic system has been associated with movement disorders (Hening et al., 2004; Kordower et al., 2013; Shetty et al., 2019), the modulation of spinal locomotor circuits (Svensson et al., 2003; Miles and Sillar, 2011; Clemens et al., 2012; Sharples et al., 2014), Willis-Ekbom disease/restless legs syndrome (Ondo et al., 2000; Clemens et al., 2006), and motor skill learning (Krakauer and Mazzoni, 2011; Wolpert et al., 2011). Most studies on the dopaminergic system demonstrate that dopaminergic neurons located in the midbrain and ventral tegmental area, defined as anatomical groups A8-10 in mammals, are functionally altered in Parkinson’s disease and addiction (Björklund and Dunnett, 2007). Interestingly, the diencephalic A11 group of dopaminergic neurons provides the sole source of spinal dopaminergic innervation in mammals (Skagerberg and Lindvall, 1985).

In zebrafish, far-projecting diencephalic dopaminergic neurons (DDNs), homologous to the A11 group in mammals, are located in the ventral diencephalic posterior tubercular (PT) area, which is anatomically delineated into the anterior rostral (PTar) and anterior caudal (PTac) groups (Ryu et al., 2007; Tay et al., 2011). In mature larvae, spontaneous DDN activity, monitored with *in vivo* calcium imaging, is synchronized both within and between PTar and PTac groups, and is correlated with locomotor and mechanosensory activity (Reinig et al., 2017). Further, PTar DDNs in mature larvae are tonically active at rest and have a phasic bursting activity pattern during locomotor activity (Jay et al., 2015). However, it is not known if activity in the DDNs changes during development.

The DDNs project to the spinal cord forming the dopaminergic diencephalospinal tract (DDT), which is considered the most conserved part of the dopaminergic system (Takada et al., 1988; Ryu et al., 2007; Kastenhuber et al., 2010). The DDT is developmentally defined in both zebrafish and mouse by the homeodomain transcription factors, Orthopedia 1a and 1b (Otpa and Otpb) (Ryu et al., 2007). The DDT is present in the spinal cord as early as 1 dpf and projects along the rostrocaudal extent of the cord by 3 dpf (Kastenhuber et al., 2010). Interestingly, the presence of dopamine transporter and tyrosine hydroxylase in the DDNs at 3 dpf implies that dopamine is available for release (Xi et al., 2011). Therefore, coarse swimming produced by immature larvae (*e.g.,* long duration episodes) could be due, in part, to a lack of functional D2-likeRs present in locomotor-related spinal neurons, a lack of endogenous dopamine release in the spinal cord, a lack of activity in the DDNs, or a combination of these. The work presented here provides evidence that dopamine receptors are present and functional in spinal circuits of immature larvae, and that increases in D2-likeR transcript levels, dopamine level, and DDN activity, correlate with the developmental transformation of locomotor activity.

## Materials and Methods

### Zebrafish Lines and Care

The University of Minnesota Institutional Animal Care and Use Committee approved all protocols used in this study. Adult zebrafish diet and recirculating system water values were described previously (Wahlstrom-Helgren et al., 2019). Experimental animals were housed in a lighted (14-hour light: 10-hour dark) 28.5°C incubator in water containing 60µg/ml Instant Ocean Sea Salt (Instant Ocean) until 7 dpf. Wildtype (WT, Segrest Farms) larvae (3-7dpf) were used in all electrophysiology, qRT-PCR, and metabolomic experiments. Double transgenic *Tg(th:Gal4^m1233^;UAS:GCaMP6s^nk13a^)* larvae (3-5 dpf) in the Casper (*roy^-/-^ nacre^-/-^)* background were used in all calcium imaging experiments.

### Spinal Transections

Larvae were anesthetized and spinal transections were made with a fine razorblade (FA-10 Feather S, Ted Pella) held with a blade breaker and holder (10053-09, Fine Science Tools). Spinalized preparations were generated in WT at 3 dpf by transecting the nervous system between body segments 3 and 4, just caudal to the hindbrain–spinal cord junction. Spinalization completely separated the brain from the spinal cord, ensuring that all descending inputs to the transected spinal cord were eliminated.

### NMDA-Induced Fictive Swimming - Peripheral Nerve Recording

Established procedures for peripheral nerve recordings were used (Masino and Fetcho, 2005; Lambert et al., 2012). Briefly, larvae were anesthetized with 0.02% Tricaine-S in extracellular saline composed of (in mM): 134 NaCl, 2.9 KCl, 1.2 MgCl_2_, 2.1 CaCl_2_, and 10 HEPES, adjusted to pH 7.8 with NaOH (10 N) and adjusted to 290-300 mOsm/L with sucrose. Larvae were pinned laterally through the notochord to a dissecting dish and subsequently spinalized. Next, the skin was removed with forceps to expose peripheral nerves, and α-bungarotoxin (25 µM in extracellular saline; 2133, Tocris) was applied for 10 min to induce paralysis. Dissecting dishes were placed on a compound microscope (BX51 WI, Olympus) and the preparations were perfused with extracellular saline that contained NMDA (50 µM) at a rate of 0.4 ml/min in a volume of approximately 1-2 ml for ∼60 min to elicit NMDA-induced fictive swimming (Fig. 1). Glass suction electrodes were positioned on the peripheral nerves at intermyotomal clefts, and recordings were initiated after 10-15 min of perfusion. Voltage signals were amplified with an Axon Multiclamp 700B amplifier connected to a Digidata 1440A digitizer. Signals were sampled at 10 kHz, band-pass filtered to 100-1000 Hz, and recorded using pClamp 10 software (Molecular Devices).

**Figure 1.**
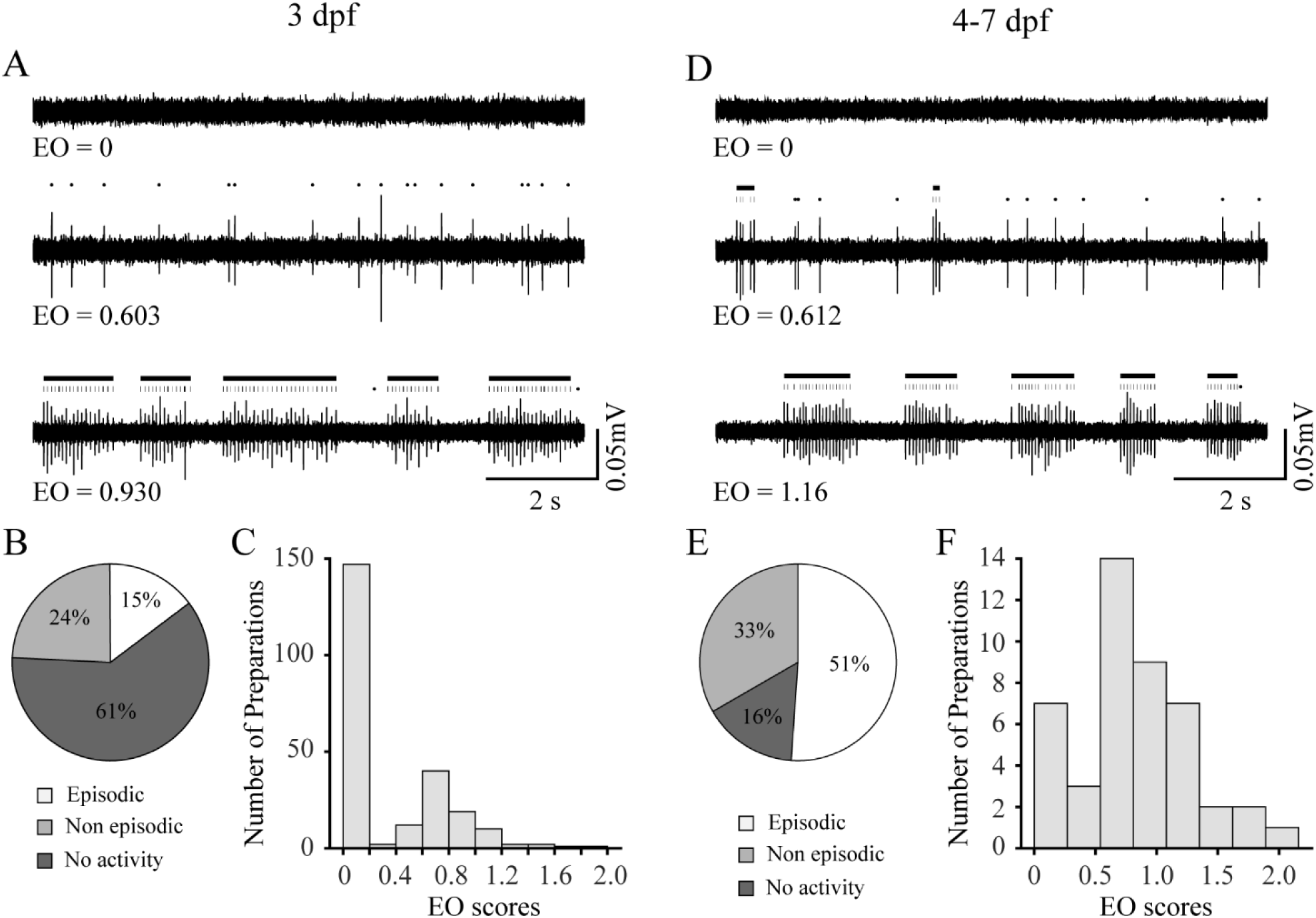
Episodically organized NMDA-induced fictive swimming is less prevalent in immature larvae. (**A**) Representative peripheral nerve recordings of fictive locomotor activity from immature (3 dpf) larvae demonstrate three outcomes: no activity (top trace), non-episodic activity (middle trace), and episodic activity (bottom trace). Black dots above the middle and bottom traces indicate peripheral nerve bursts not organized into episodes, while vertical black lines indicate peripheral nerve bursts organized into episodes. Black horizontal bars indicate episodes. Values for episodic organization (EO) are indicated below the traces. (**B**) Pie chart showing the proportion (15% (35 of 236); white) of immature larvae generate episodically organized (EO ≥ 0.75) locomotor activity. (**C**) Frequency histogram showing the distribution of EO scores for immature larvae. (**D**) Representative peripheral nerve recordings of fictive locomotor activity from mature (4-7 dpf) larvae demonstrate three outcomes: no activity (top trace), non-episodic activity (middle trace), and episodic activity (bottom trace). Black dots above the middle and bottom traces indicate peripheral nerve bursts not organized into episodes, while vertical black lines indicate peripheral nerve bursts organized into episodes. Black horizontal bars indicate episodes. Values for episodic organization (EO) are indicated below the traces. (**E**) Pie chart showing the proportion (51% (23 of 45); white) of mature larvae generate episodically organized (EO ≥ 0.75) locomotor activity. (**F**) Frequency histogram showing the distribution of EO scores for mature larvae.

Peripheral nerve recordings were analyzed by an in-house program written in MATLAB (The MathWorks) to measure properties of fictive swimming described previously (Wiggin et al., 2012). Briefly, baseline NMDA-induced locomotor properties were measured first during a two-minute analysis window following a 30 min washout of anesthetic and paralytic. A second measurement (two-minute window) was repeated after 10 min application of vehicle or drug (DMSO, D2-likeR agonist, or D4R agonist respectively). Episodic organization (EO) scores were calculated at baseline, as described previously (Wiggin et al., 2012; Montgomery et al., 2021). Briefly, EO was measured as the log_10_ ratio of the means of long inter-burst periods to short inter-burst periods. Short (inter-episode-like) and long (intra-episode-like) inter-burst periods were determined using a critical value of two standard deviations above the mean of the burst periods for each preparation. Higher EO scores represent a large difference between these means, such that bursts are organized into discrete episodes. Lower EO scores represent a small difference between these means, such that burst are less organized into discrete episodes. An EO score of zero (0) represented experimental preparations that either generated no locomotor activity or that had < 50 total bursts within the 2-minute analysis window. The analysis of pharmacological perturbation experiments was restricted to preparations that produced an EO score of ≥ 0.75 (Fig. 1) and that had > 200 bursts within the 2-minute analysis window. Episodes were defined as collections of at least three peripheral nerve bursts induced by NMDA separated by less than 150 msec of quiescence between bursts (Wiggin et al., 2012). Total burst number was defined as the count of peripheral nerve bursts recorded during the 2-minute analysis window. Episode duration was defined as the time of onset of the first burst of an episode until the offset of the final burst of the same episode. All the episode durations within the two-minute analysis window were averaged to find the mean episode duration for each timepoint.

### Pharmacology

Stock solutions of NMDA (10 mM; M3262, Sigma Aldrich) were dissolved in extracellular saline and diluted in extracellular saline to working concentration (50 µM) Stock solutions of the D2-likeR-specific agonist (quinpirole (3 mM); Q102, Sigma Aldrich) and D4R-specific agonist (PD168,077 (7.5 mM); P233, Sigma Aldrich) were dissolved in deionized water or dimethylsulfoxide (DMSO; D5879, Sigma Aldrich), respectively. Stock solutions were diluted in extracellular saline to the working concentration (quinpirole, 10 µM; PD168,077, 1 µM; DMSO, 0.02%) determined previously (Lambert et al., 2012).

### Reverse Transcription Quantitative Polymerase Chain Reaction

Larvae from four separate clutches (groups) were collected at 3 and 5 dpf. To separate the heads from the bodies, larvae were anesthetized (0.04% Tricaine, MS-222), placed on a bed of Sylgard, and transections were performed as described above. The transected tissue was then placed immediately in saline on ice. The heads and bodies from 200 larvae/group were processed separately for the extraction of total RNA (RNeasy Mini Kit, Qiagen) using routine procedures (Martin et al., 2019; Parvin et al., 2021).

Total RNA (100 ng) was reverse transcribed to produce cDNA using TaqMan EZ RT-PCR Core reagents (Applied Biosystems). The expression levels of D2-like receptor transcripts (*drd2a, drd2b, drd2l, drd3, drd4a, drd4b, drd4rs*) were measured by TaqMan quantitative real-time (qRT)-PCR using a 7900HT Fast Real-Time PCR System (Applied Biosystems), as replicates (x2). *β-actin* served as a reference gene. Validated qPCR assays (probes, forward and reverse primers) were used for the following genes: *drd2a* (Assay ID Dr03106158_m1), *drd2b* (Assay ID Dr03093765_m1), *drd2l* (Assay ID Dr03119255_m1), *drd3* (Assay ID Dr03131905_m1), *drd4a* (Assay ID Dr03090314_m1), *drd4b* (Assay ID Dr03096340_m1), *drd4-rs* (Assay ID Dr03096342_m1), and *actin* (Assay ID Dr03432610_m1). All assays were obtained from ThermoFisher Scientific. The targeted gene accession numbers from NCBI are: *drd2a* (NM_183068.1), *drd2b* (NM_197936.1), *drd2l* (NM_197935.1), *drd3* (Dr03131905_m1), *drd4a* (NM_001012616.3), *drd4b* (NM_001012618.1), *drd4-rs* (NM_001012620.3), and *actin* (NM_131031.2). The delta-delta Ct method (2^(Δ-Δ CT)) was used to measure relative changes in gene expression. The expression for each gene of interest was normalized to *β-actin* expression and presented as a fold change, increase or decrease, in 3 dpf groups, which were normalized to 1, and compared to 5 dpf groups.

### Liquid Chromatography-Mass Spectrometry

Samples of intact zebrafish were collected from three separate clutches on days 3 and 5 post-fertilization. Each of the samples was reconstituted in lysis buffer (210 µL; 97.8% water, 2% acetonitrile, and 0.2% formic acid) and ^13^C_6_-dopamine (2 ng; CLM-3369-PK, Cambridge Isotope Laboratories, Inc.) and transferred to Precellys homogenization tubes (500 µL; 432-0293, Avantor). Next, samples were subjected to three, 30 second pulses of homogenization at 5800 revolutions per minute (rpm) in a Precellys Evolution homogenizer (P002511-PEVT0-A.0, Betin Corp.). Following homogenization, supernatant (10 µL) was removed from each sample and used to quantify protein via Bradford Assay (23200, Thermo Fisher Scientific) to normalize dopamine content to total protein. To the remaining samples, methanol (200 µL) was added and the homogenization repeated in the Precellys homogenizer using the same settings. Following the second homogenization, samples were centrifuged at 13,200 rpm for 15 minutes at room temperature. The supernatant from each sample was then transferred to a 1.7 ml microcentrifuge tube (022363204, Eppendorf) after which chloroform (800 µL) was added to samples. The samples were vortexed briefly, shaken at 600 rpm, incubated at room temperature for 10 minutes, and centrifuged again for 15 minutes at 13,200 rpm at room temperature. The aqueous upper layer of each sample was removed and transferred to a new 1.7 ml microcentrifuge tube (022363204, Eppendorf) and dried overnight in a SpeedVac SPD210 (Thermo Fisher Scientific). Samples were then reconstituted in water (90 µL) with 0.1% formic acid for analysis.

For dopamine (36532, Cayman Chemical) quantitation, a calibration curve was constructed with dopamine concentrations of 0, 0.1, 0.5, 1, 5, 10, 50, and 100 ng/mL in water with 0.1% formic acid and a constant ^13^C_6_-dopamine (Cambridge Isotope Laboratories, Inc., Tewksbury, MA) concentration of 100 ng/mL. Next, samples were analyzed on a mass spectrometer (Sciex QTrap 6500+, SCIEX) in positive mode interfaced with a Waters Acquity UPLC plumbed with a Waters Acquity UPLC BEH C18 Column (130Å, 1.7 µm, 2.1 mm X 50 mm; Waters Corp). Mass spectrometer method settings included a curtain gas flow of 25 pounds per square inch (psi), a collision gas value of medium, an ion spray voltage of 5500 V, a temperature of 350°C, an ion source gas 1 value of 20 psi and an ion source gas 2 value of 30 psi. Mobile phase A consisted of water with 0.1% formic acid, while mobile phase B consisted of acetonitrile with 0.1% formic acid. Samples were run on a 15-minute method at 0.2 mL/min with a solvent gradient consisting of 98% A from 0 to 0.1 minutes, 98% to 65% A from 0.1 to 5 minutes, 65% to 2% A from 5 to 11 minutes, 2% A from 11 to 12 minutes, 2% to 98% A from 12 to 13.5 minutes, and 98% A from 13.5 to 15 minutes. Each sample was run four times with fresh aliquots (20 µL) injected for each run. Raw data was analyzed using the Skyline suite (https://skyline.ms/project/home/begin.view).

### In Vivo Calcium Imaging – longitudinal assay

Transgenic zebrafish larvae (*th:Gal4^m1233^;UAS:GCaMP6s^nk13a^*) were anesthetized with 0.02% Tricaine-S (Western Chemical) in extracellular saline and embedded (positioned dorsal ventrally) in 0.2% low melting point agarose. Dissecting dishes were placed on an (BX51WI, Olympus) compound microscope and perfused with extracellular saline at a rate of 0.4 ml/min in a volume of approximately 1-2 ml with recordings initiated after 20-30 min of perfusion. GCaMP signal was visualized using epifluorescent blue light (1.9 mW/cm^2^; X-Cite 12-LED Boost lamp, Excelitas Technologies) and 20x/NA 0.5 water immersion objective (UMPlanFL N, Olympus) and a fluorescence imaging camera (Retiga EXi, QImaging). Calcium signals were collected at 5 Hz for 30 minutes. Subsequently, each larva was unembedded and returned to embryo water and *in vivo* calcium imaging was repeated at 3 and 5 dpf.

An anatomical atlas was used as a guide to identify bilateral DDN/PTac neurons (Haehnel-Taguchi et al., 2018). Bilateral ROIs of equal size were placed over regions of identified DDN/PTac neurons (Fig 8A & D). To control for background signal, an equal sized ROI was placed over a region devoid of indicator and neurons. Calcium signals were bleach corrected using Fiji (Schindelin et al., 2012) and changes in fluorescence (ΔF/F) were measured using Microsoft Excel (Microsoft Corporation). The ‘findpeaks’ function in MATLAB was used to find the number of peaks and the magnitude of each peak. The minimum interval between peaks was set to 5 frames (1 sec) and the minimum ΔF/F prominence was set to 0.35. To determine the average power of the calcium signals for each recording, the root mean square of the signals (ΔF/F) was determined and squared.

### Statistical Analysis

All statistical analyses were performed with SigmaPlot 16.0 (Graffiti). Episodic properties of locomotor activity, measured with peripheral nerve recordings, were analyzed using Student’s t-test or paired t-tests for parametric data and Mann-Whitney Rank Sum test or Wilcoxon Signed Rank test for non-parametric data. Calcium transients recorded from bilateral DDN/PTac neuron groups were analyzed using Pearson’s Product Moment Correlation and Student’s t-tests for parametric data and Mann-Whitney Rank Sum test for non-parametric data. qRT-PCR of D2-likeR mRNA expression between 3 and 5 dpf larvae were analyzed using Student’s t-test with Prism v9 (GraphPad Software). Metabolomic analysis of DA levels were analyzed using Students t-test (Origin 2025 Software). All experimenters were blinded to age for qRT-PCR and metabolomic analysis. Significance was established using an α criterion of *p* = 0.05. In the figures, \**p* < 0.05, \*\**p* < 0.01, and \*\*\**p* < 0.001. Data are expressed as means with standard deviation (mean (SD)).

## Results

### The spinal locomotor circuit is less functionally organized in immature larvae

Spontaneous swimming episodes produced by immature larvae are less frequent and of longer duration than those produced by mature larvae, indicating that locomotor activity transforms during larval development (Lambert et al., 2012). To assess potential differences in function of the spinal locomotor circuit before and after the transformation of locomotor activity, we compared, in spinalized preparations, NMDA-induced locomotor properties between immature and mature larvae (Fig. 1). The proportion of immature larvae that did not generate NMDA-induced activity (Fig. 1A, top trace & B) was significantly greater (Table 1) compared to the proportion of mature larvae (Fig. 1D, top trace & E). In addition, the proportion of immature larvae that produced episodically organized locomotor activity (Fig. 1A, bottom trace & B) was significantly lower (Table 1) compared to mature larvae (Fig. 1D, top trace & E).

**Table 1.** Proportions of immature and mature larvae that produce NMDA-induced locomotor responses.

|  | Group |  |
| --- | --- | --- |
|  | 3 dpf | 4-7 dpf |
| No activity (%) | 61.0<br>(144 of 236) | 15.6<br>(7 of 45) |
|  | z = 5.61<br>p < 0.001 |  |
| Non-episodic (%) | 24.2<br>(57 of 236) | 33.3<br>(15 of 45) |
|  | z = 1.29<br>p = 0.196 |  |
| Episodic (%) | 14.8<br>(35 of 236) | 51.1<br>(23 of 45) |
|  | z = 5.51<br>p < 0.001 |  |

Next, we performed an initial analysis which included all preparations that generated locomotor activity with an EO score greater than zero (0) (Figs. 1A & D, middle and bottom traces). Neither episode durations nor the number of bursts generated by immature larvae were significantly different (Table 2) compared to mature larvae. However, the EO scores produced by immature larvae were significantly lower (Table 2) compared to mature larvae.

**Table 2.**
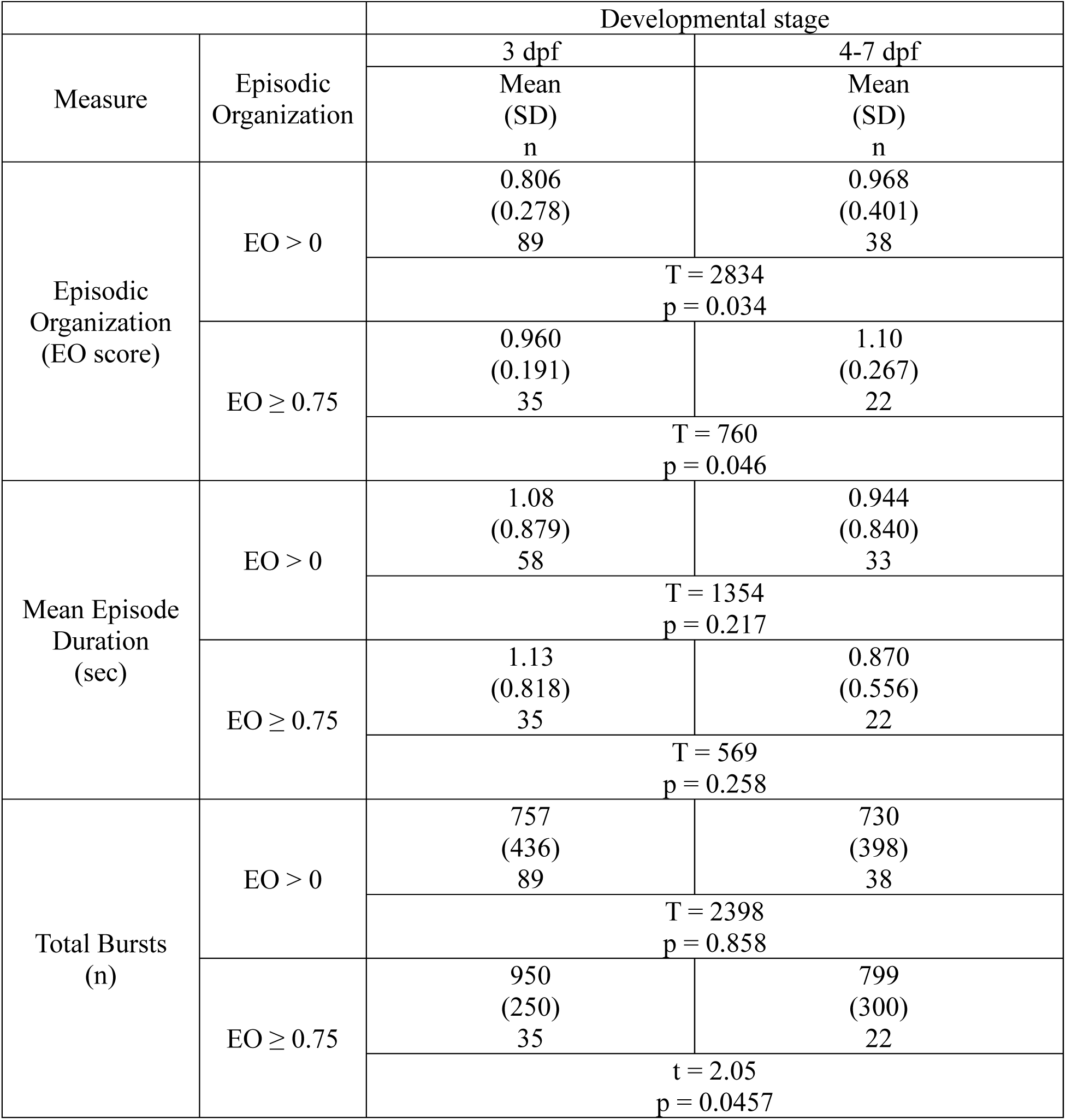
Comparison of NMDA-induced locomotor activity measures between immature and mature larvae. Measures reported as mean (SD).

Finally, we restricted our analysis to only preparations that produced NMDA-induced locomotor activity with an EO score greater than 0.75 (Fig. 1 A & D, bottom traces). Although episodes generated by immature larvae were well organized, the EO scores were significantly lower, and the total number of bursts were significantly higher (Table 2) compared to mature larvae. However, episode durations generated by immature larvae were not significantly different (Table 2) compared to mature larvae. Although the spinal locomotor circuit in immature larvae was less functionally organized compared to mature larvae, our results demonstrated that it was sufficiently functional in, at least, a proportion of immature larvae to produce episodically organized locomotor activity.

### D4Rs are present and functional in immature larvae

Our previous work demonstrated that dopaminergic signaling through inhibitory D2-likeRs was necessary to maintain the mature locomotor phenotype (short duration episodes) following the developmental transformation of locomotor activity (Lambert et al., 2012). Thus, we reasoned that the coarse locomotor pattern produced by immature larvae could be due to a lack of functional D2-likeRs present in locomotor-related spinal neurons.

First, we assessed potential time-dependent (early vs. late) effects on the episodic properties of NMDA-induced locomotor activity in immature larvae (Fig 2). During continuous NMDA application, significant differences between early and late conditions were not found for mean episode duration, EO score, or total number of bursts (Fig. 2B-D; Table 3). These results demonstrated that NMDA-induced activity remained episodically organized and stable for the duration (∼60 min) of the experiment.

**Figure 2.**
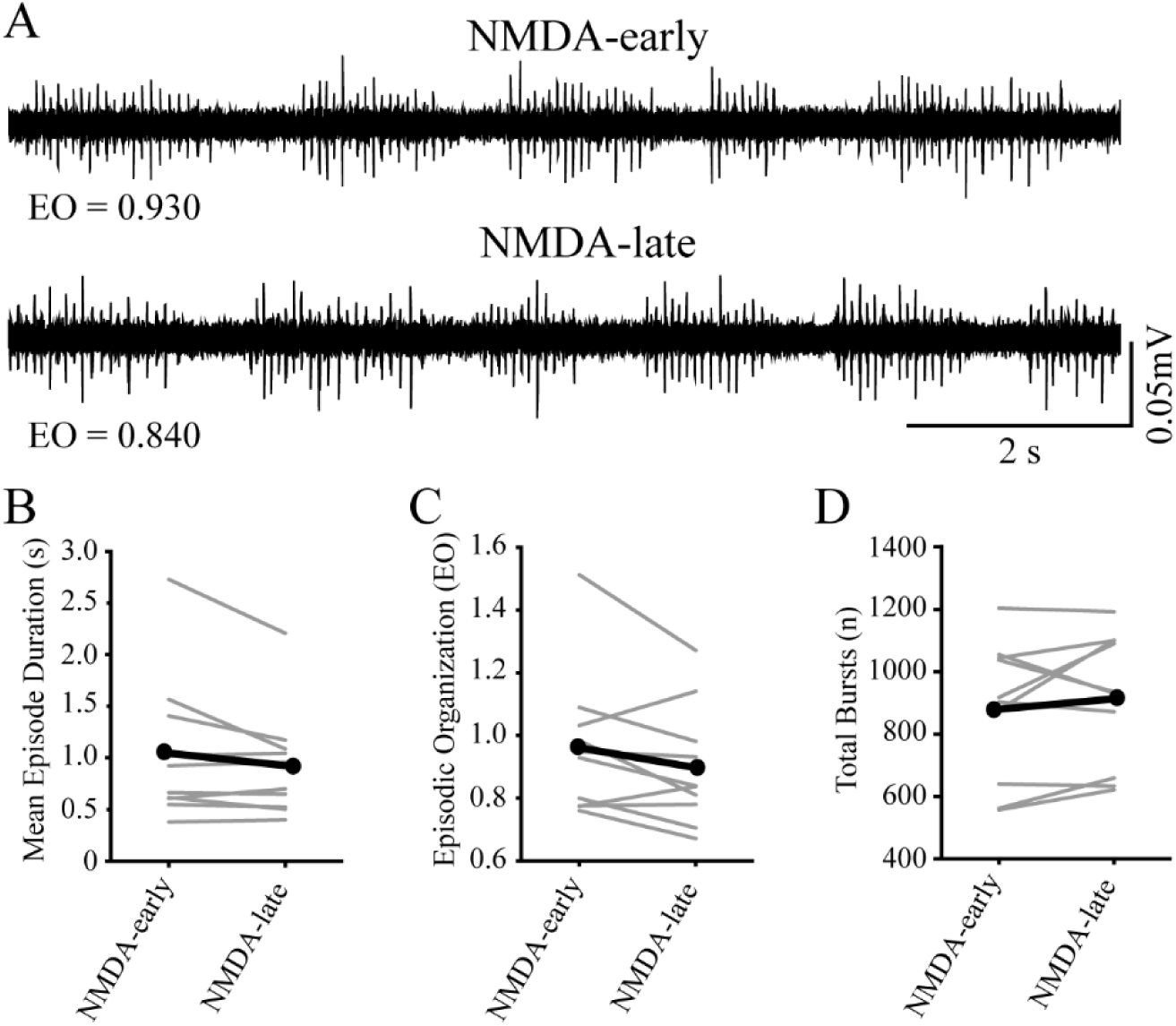
NMDA-induced spinal locomotor activity remains episodically organized and stable. (**A**) Representative peripheral nerve recordings from an individual preparation acquired at early (NMDA-early, 30 min) and late (NMDA-late, 40 min) timepoints during continuous NMDA application. Values for episodic organization (EO) are indicated below the traces. (**B**) Episode duration is not different between NMDA-early and NMDA-late timepoints. Gray lines indicate within subjects repeated measures, while black lines represent the mean (n = 10 larvae). (**C**) Episodic organization is not different between NMDA-early and NMDA-late timepoints. Gray lines indicate within subjects repeated measures, while black lines represent the mean (n = 10 larvae). (**D**) The total number of bursts are not different between NMDA-early and NMDA-late timepoints. Gray lines indicate within subjects repeated measures, while black lines represent the mean (n = 10 larvae).

**Table 3.** Comparison of locomotor properties in immature larvae under different experimental conditions. Measures are reported as mean (SD).

| Experimental Condition | Mean Episode Duration (sec) |  | Episodic Organization (EO score) |  | Total Bursts (n) |  |
| --- | --- | --- | --- | --- | --- | --- |
|  | Baseline Mean (SD) | Treatment Mean (SD) | Baseline Mean (SD) | Treatment Mean (SD) | Baseline Mean (SD) | Treatment Mean (SD) |
| Control | 1.04<br>(0.705) | 0.924<br>(0.526) | 0.961<br>(0.227) | 0.898<br>(0.190) | 880<br>(224) | 913<br>(214) |
|  | Z = -1.07<br>p = 0.322<br>n = 10 |  | t = 1.90<br>p = 0.0900<br>n = 10 |  | t = -0.951<br>p = 0.367<br>n = 10 |  |
| D2-likeR agonist | 0.936<br>(1.14) | 0.523<br>(0.582) | 0.841<br>(0.0722) | 0.645<br>(0.110) | 800<br>(282) | 695<br>(372) |
|  | Z = -2.02<br>p = 0.063<br>n = 5 |  | t = 3.98<br>p = 0.0164<br>n = 5 |  | t = 1.16<br>p = 0.312<br>n = 5 |  |
| D4R agonist | 2.03<br>(0.651) | 1.46<br>(0.604) | 1.11<br>(0.134) | 0.894<br>(0.123) | 1234<br>(129) | 1219<br>(165) |
|  | t = 3.14<br>p = 0.0202<br>n = 7 |  | t = 5.27<br>p = 0.00188<br>n = 7 |  | t = 0.448<br>p = 0.670<br>n = 7 |  |
| DMSO | 0.668<br>(0.540) | 0.479<br>(0.356) | 0.937<br>(0.105) | 0.813<br>(0.0683) | 817<br>(263) | 840<br>(284) |
|  | Z = -2.02<br>p = 0.063<br>n = 5 |  | t = 2.07<br>p = 0.107<br>n = 5 |  | t = -0.506<br>p = 0.639<br>n = 5 |  |

Next, to establish that inhibitory dopamine receptors were present and functional in immature larvae, we compared episodic properties of NMDA-induced locomotor activity before (baseline) and during application (treatment) of D2-likeR (quinpirole) or D4R-specific (PD168,077)) agonists. Application of the D2-likeR agonist produced a non-significant decrease in episode durations (Fig. 3A & B; Table 3), a significant decrease in EO score (Fig. 3A & C; Table 3), and no effect on the number of bursts (Fig. 3A & D; Table 3). Application of the D4R agonist produced significant decreases for both episode durations (Fig. 4A & B; Table 3) and EO scores (Fig. 4A & C; Table 3), but no effect on the number of bursts (Fig. 4A & D; Table 3).

**Figure 3.**
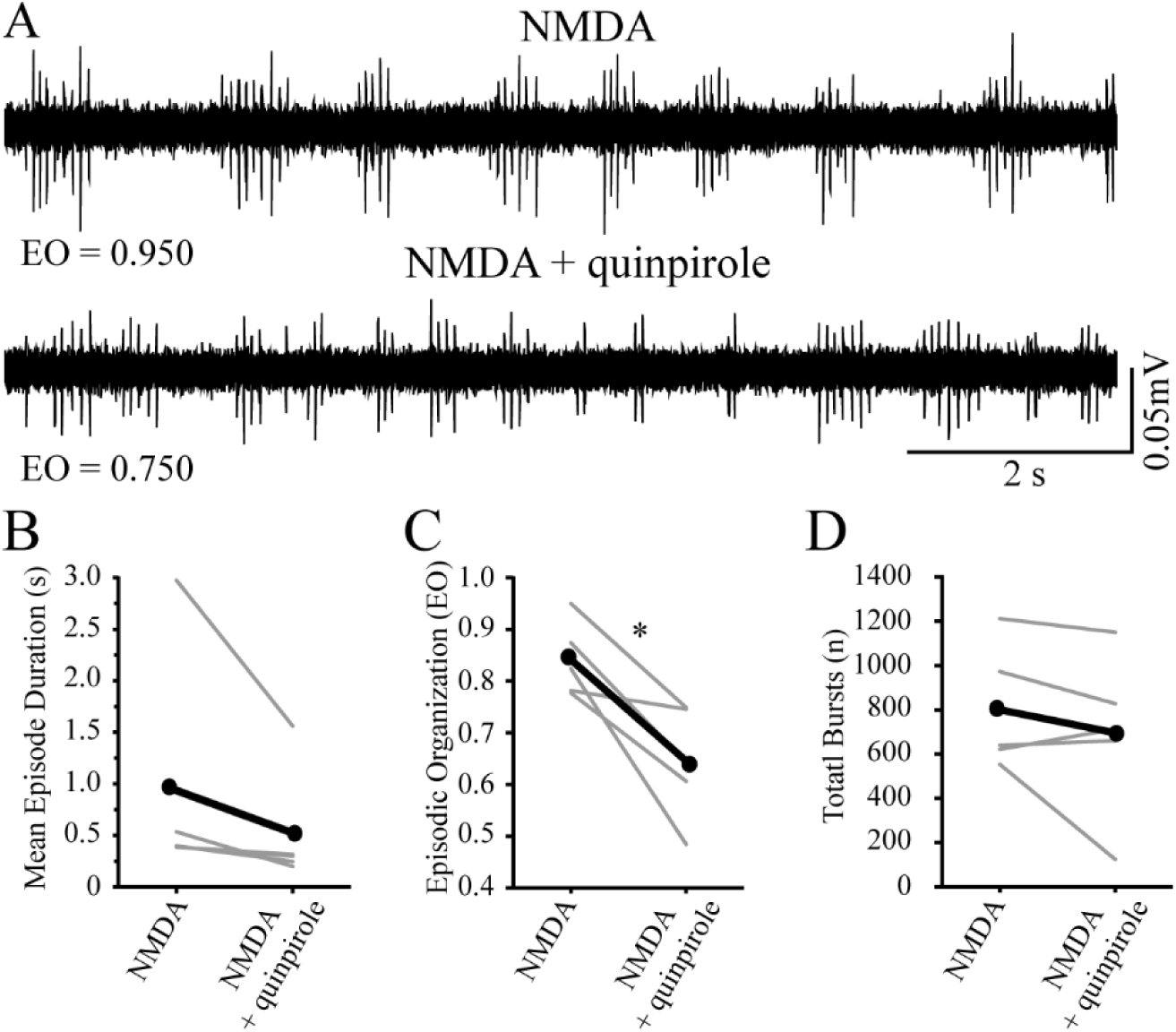
Application of exogenous D2-like receptor subtype agonist during NMDA-induced fictive swimming in immature larvae decreases episodic organization. (**A**) Representative peripheral nerve recordings from an individual preparation acquired at baseline (NMDA) and during agonist treatment (NMDA + quinpirole). Values for episodic organization (EO) are indicated below the traces. (**B**) Episode duration decreases, but not significantly, during treatment (NMDA + quinpirole) compared to baseline (NMDA). Gray lines indicate within subjects repeated measures, while black lines represent the mean (n = 5 larvae). (**C**) Episodic organization decreases during treatment (NMDA + quinpirole) compared to baseline (NMDA). Gray lines indicate within subjects repeated measures, while black lines represent the mean (n = 5 larvae). (**D**) The total number of bursts are not affected during treatment (NMDA + quinpirole) compared to baseline (NMDA). Gray lines indicate within subjects repeated measures, while black lines represent the mean (n = 5 larvae). Asterisk indicates significant differences at p < 0.05.

**Figure 4.**
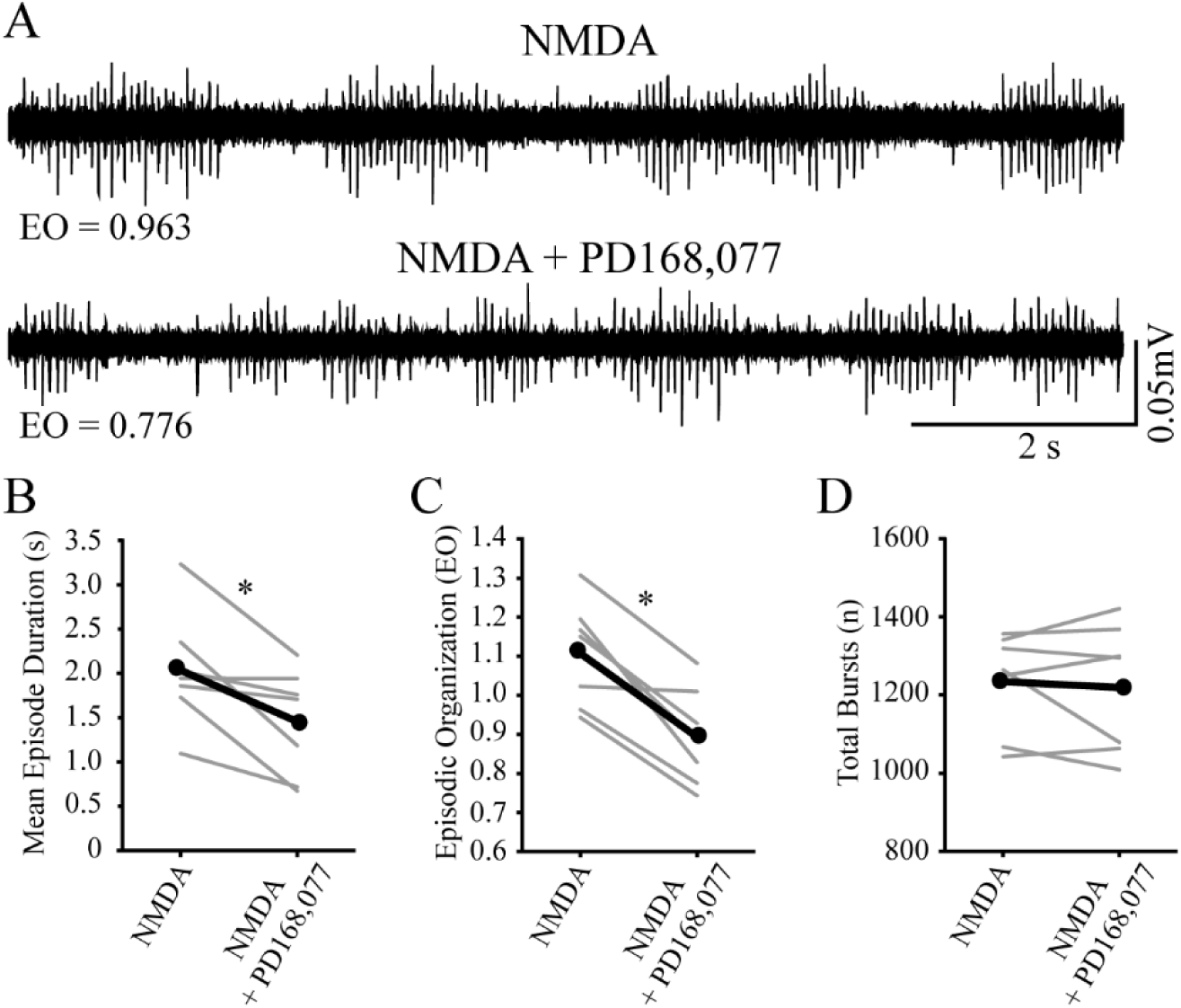
Application of exogenous D4-specific receptor subtype agonist during NMDA-induced fictive swimming in immature larvae decreases episode duration and episodic organization. (**A**) Representative peripheral nerve recordings from an individual preparation acquired at baseline (NMDA) and during agonist treatment (NMDA + PD 168,077). Values for episodic organization (EO) are indicated below the traces. (**B**) Episode duration decreases during treatment (NMDA + PD 168,077) compared to baseline (NMDA). Gray lines indicate within subjects repeated measures, while black lines represent the mean (n = 7 larvae). (**C**) Episodic organization decreases during treatment (NMDA + PD 168,077) compared to baseline (NMDA). Gray lines indicate within subjects repeated measures, while black lines represent the mean (n = 7 larvae). (**D**) The total number of bursts are not affected during treatment (NMDA + PD 168,077) compared to baseline (NMDA). Gray lines indicate within subjects repeated measures, while black lines represent the mean (n = 7 larvae). Asterisks indicate significant differences at p < 0.05.

Finally, since DMSO was used as the vehicle for the D4R-specific agonist, we assessed potential effects of vehicle administration on the episodic properties of NMDA-induced locomotor activity. There were no significant differences in the mean duration of episodes (Fig. 5A & B; Table 3), EO score (Fig. 5A & C; Table 3), or number of bursts (Fig. 5A & D; Table 3) during application of DMSO. These results demonstrated that, minimally, D4Rs were present and functional in the spinal locomotor circuits of immature larvae.

**Figure 5.**
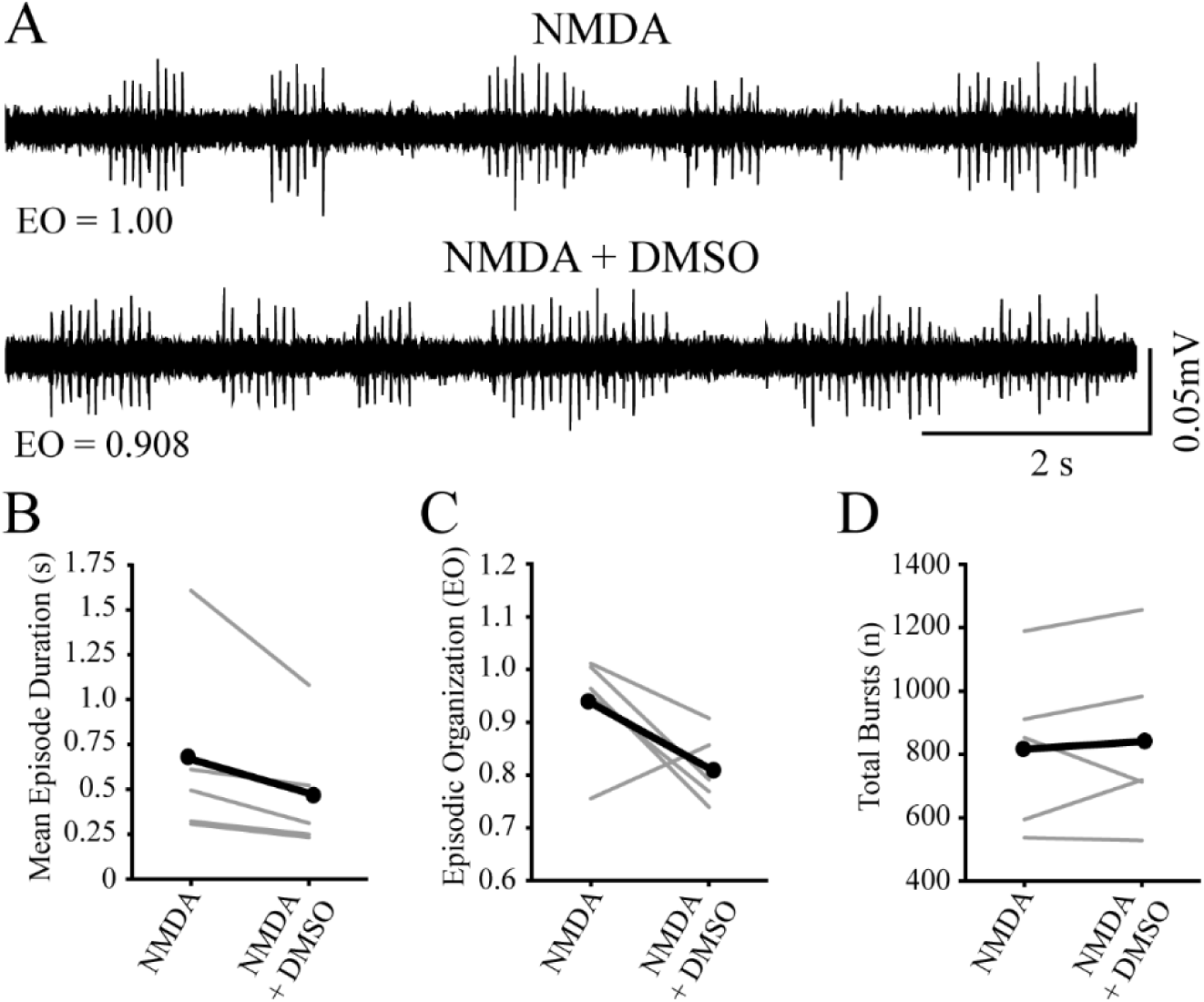
Application of DMSO during NMDA-induced fictive swimming in immature larvae does not affect episode duration, episodic organization, or total number of bursts. (**A**) Representative peripheral nerve recordings from an individual preparation acquired at baseline (NMDA) and during treatment (NMDA + DMSO). Values for episodic organization (EO) are indicated below the traces. (**B**) Episode duration is not affected during treatment (NMDA + DMSO) compared to baseline (NMDA). Gray lines indicate within subjects repeated measures, while black lines represent the mean (n = 5 larvae). (**C**) Episodic organization is not affected during treatment (NMDA + DMSO) compared to baseline (NMDA). Gray lines indicate within subjects repeated measures, while black lines represent the mean (n = 5 larvae). (**D**) The total number of bursts are not affected during treatment (NMDA + DMSO) compared to baseline (NMDA). Gray lines indicate within subjects repeated measures, while black lines represent the mean (n = 5 larvae).

### Gene expression of all D2-likeRs is greater in mature larvae

Several lines of evidence suggest that dopamine receptor genes are differentially expressed during development. First, the expression of dopamine receptor genes (*drd1, drd2a, drd2b, drd2l, drd3,* and *drd4*) are detected in the brain and/or spinal cord by 2 dpf, and an increase in expression during development (from 2 to 5 dpf) has been described for at least some genes (*drd1* and *drd2b*) (Boehmler et al., 2004; Li et al., 2007). Second, we previously reported that the transformation of locomotor activity during development, from long duration swim episodes produced by immature larvae to short duration swim episodes produced by mature larvae, was dependent on D4R signaling (Lambert et al., 2012). Finally, we demonstrated in this study that D4Rs were present and functional in immature larvae (Figs. 3 & 4). However, potential developmental changes in gene expression levels of all D2-likeR subtypes in the spinal cord that function to support this locomotor transformation have not been quantified.

Therefore, to compare the levels of gene expression of the D2-likeR subtypes between immature and mature larvae, qRT-PCR was performed on transected bodies as a proxy to analyze mRNA expression in spinal cord. Specifically, we assessed the expression levels of *drd2a*, *drd2b*, *drd2l, drd3*, *drd4a, drd4b,* and *drd4-rs*) mRNA. All mRNA targets, except for *drd2l*, were significantly increased in mature larvae compared to immature larvae (Fig. 6). We also assessed mRNA expression of the D2-likeR subtypes in the heads of mature and immature larvae. Consistent with the changes observed in the body, the mRNA levels of all seven D2R targets were significantly increased in mature larvae (data not shown).

**Figure 6.**
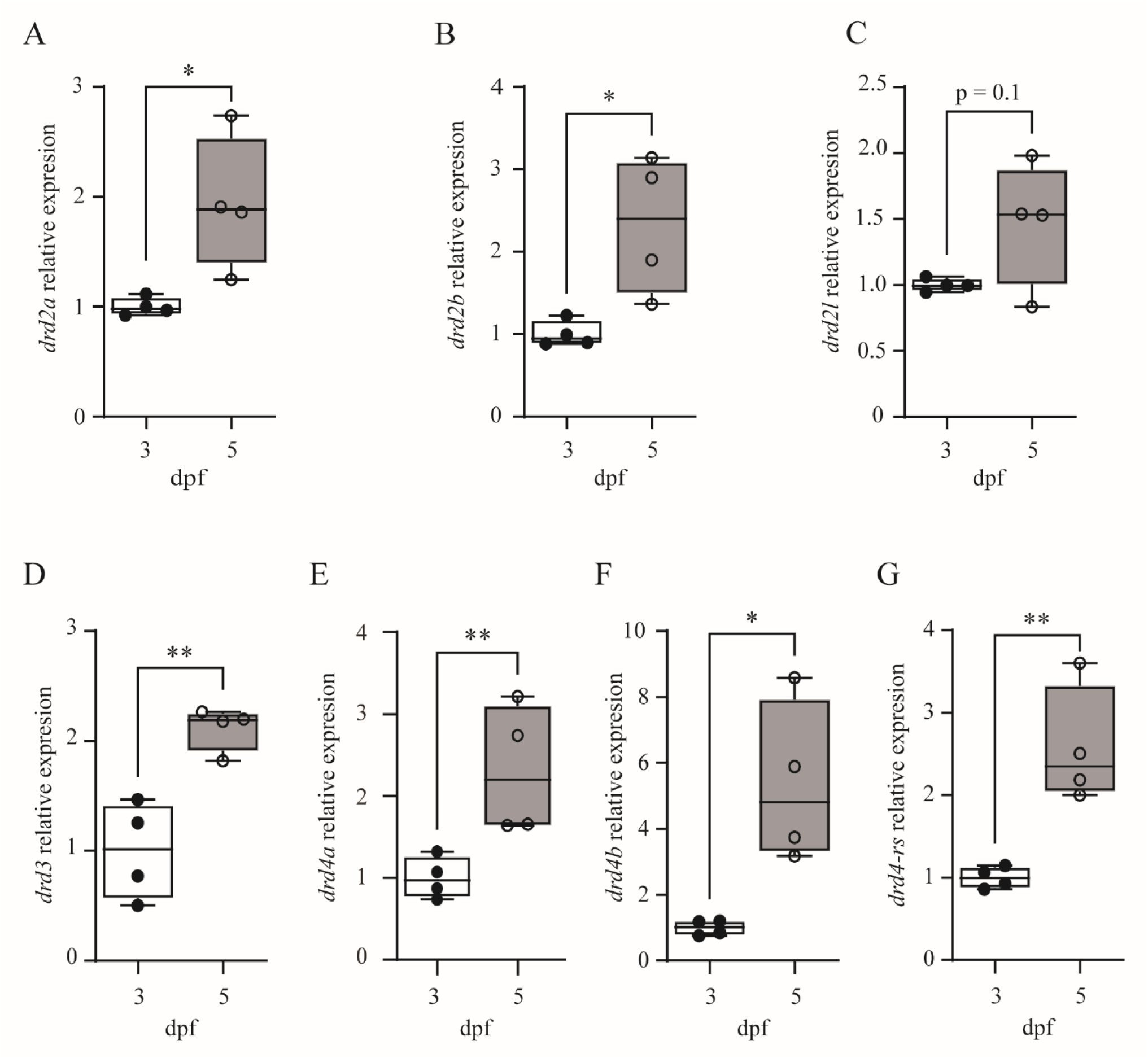
Gene expression of most D2-like receptor subtypes is greater in mature larvae. **(A-G)** Plots show that expression of all D2-like receptor subtypes, except for *drd2l*, from transected bodies of larval zebrafish are significantly increased at 5 dpf (open circles, N = 4) compared to 3pf (black-filled circles, N = 4) *drd2a,* p = 0.0228; *drd2b,* p = 0.0205; *drd2l,* p = 0.096; *drd3*, p = 0.0037; *drd4a,* p = 0.0196; *drd4b,* p = 0.0124; *drd4-rs*, p = 0.005; N = 4 at 3 dpf; N = 4 at 5 dpf)

These results demonstrated that gene expression of all D2-likeR subtypes increased in the bodies of larval zebrafish across the developmental transformation of locomotor activity (Fig. 6). This increase in gene expression in mature larvae is consistent with the hypothesis that locomotor activity is refined by an increase in dopaminergic signaling during development.

### Dopamine levels are greater in mature larvae

Our previous work demonstrated that dopaminergic signaling was required for the developmental transformation of locomotor activity (Lambert et al., 2012), that D4Rs were present and functional in the spinal locomotor circuits of immature larvae (Figs. 3 & 4), and that gene expression of all D2-likeR subtypes increased in the bodies of larval zebrafish across the developmental transformation of locomotor activity (Fig. 6). Thus, we next tested the hypothesis that an increase in DA levels correlated with the transformation of locomotor activity.

To compare absolute levels of dopamine between immature and mature larvae, we performed liquid chromatography-mass spectrometry on intact larvae at 3 and 5 dpf. Dopamine levels were significantly greater in mature larvae compared to immature larvae (Fig. 7). These results are consistent with the hypothesis that locomotor activity is transformed through an increase in dopaminergic signaling during development.

**Figure 7.**
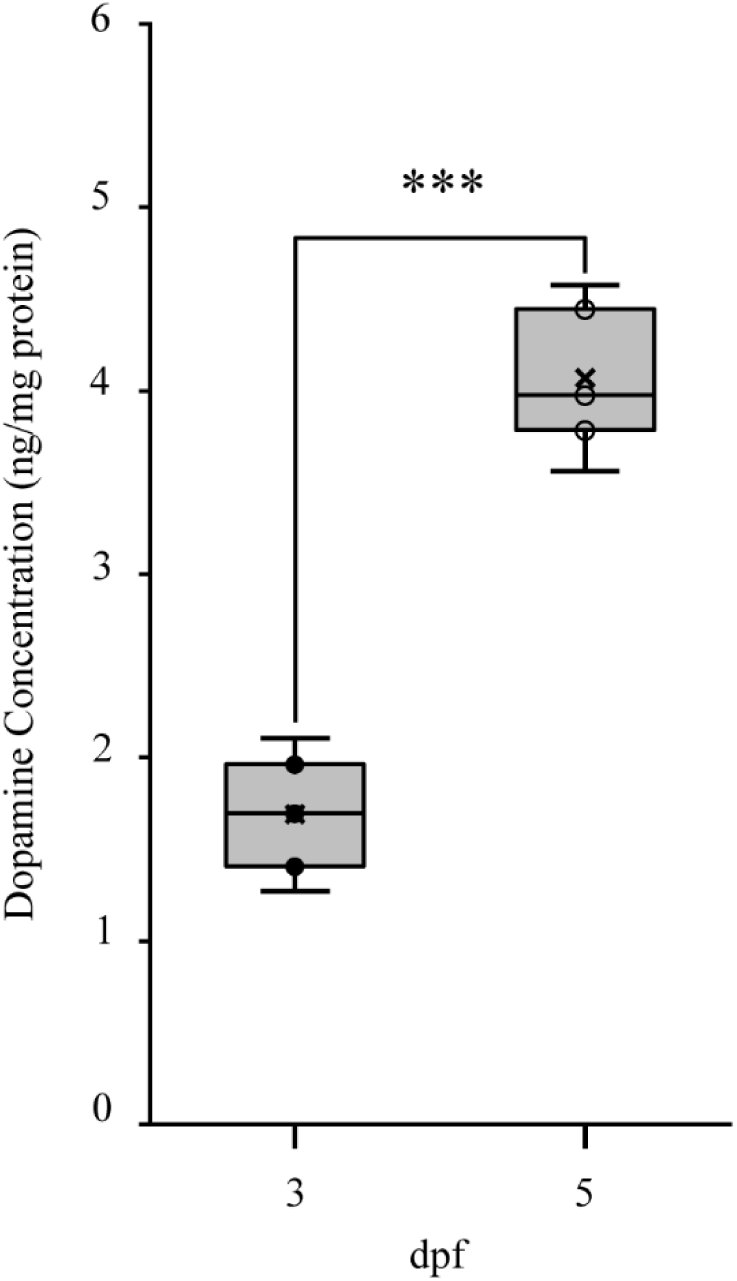
Dopamine levels are greater in mature larvae. Plot shows that the absolute amount of dopamine from intact larvae are significantly increased (t = -9.41, p = 7.10e-4, N = 3) at 5 dpf (open circles, mean (SD) = 4.07 (0.339)) compared to 3pf (black-filled circles, mean (SD) = 1.69 (0.278)). Asterisk indicates significant difference at \*\*\**p* < 0.001.

### DDN activity is greater in mature larvae

Although the level of dopamine in mature larvae is greater than in immature larvae (Fig. 7), direct evidence demonstrating that dopamine is more readily released in the spinal cord of mature larvae is lacking. We hypothesized that increased DDN/PTac activity from the immature to the mature stage of larval development was responsible for locomotor transformation. Therefore, we used DDN/PTac activity as a proxy for dopamine release. Since a larger amount of dopamine was present in mature larvae (Fig. 7), then an increase in activity in DDN/PTac neurons in mature larvae would increase the yield of dopamine release on spinal locomotor circuits.

To compare neural activity between immature and mature larvae, we measured *in vivo* calcium transients in bilateral DDN/PTac neurons within individual larvae (repeated measures) before (3 dpf) and after (5 dpf) the developmental transformation of locomotor activity. Since neural activity between the left and right groups of DDN/PTac neurons were highly correlated at both 3 dpf and 5 dpf (Fig. 8E & F; Table 4), measures of calcium signals (number, peak, magnitude) were averaged within each preparation. The number of calcium transient peaks were greater in mature larvae compared to immature larvae (Fig. 8G; Table 4). The magnitude of calcium transients was significantly greater in mature larvae compared to immature larvae (Fig. 8H; Table 4). Finally, power measures of the calcium transients were significantly greater in mature larvae compared to immature larvae (Fig. 8I; Table 4). These results demonstrated that DDN/PTac activity was greater in mature larvae, which supports the hypothesis that increased dopaminergic signaling through DDN/PTac activity correlates with the transformation of locomotor activity during development.

**Figure 8.**
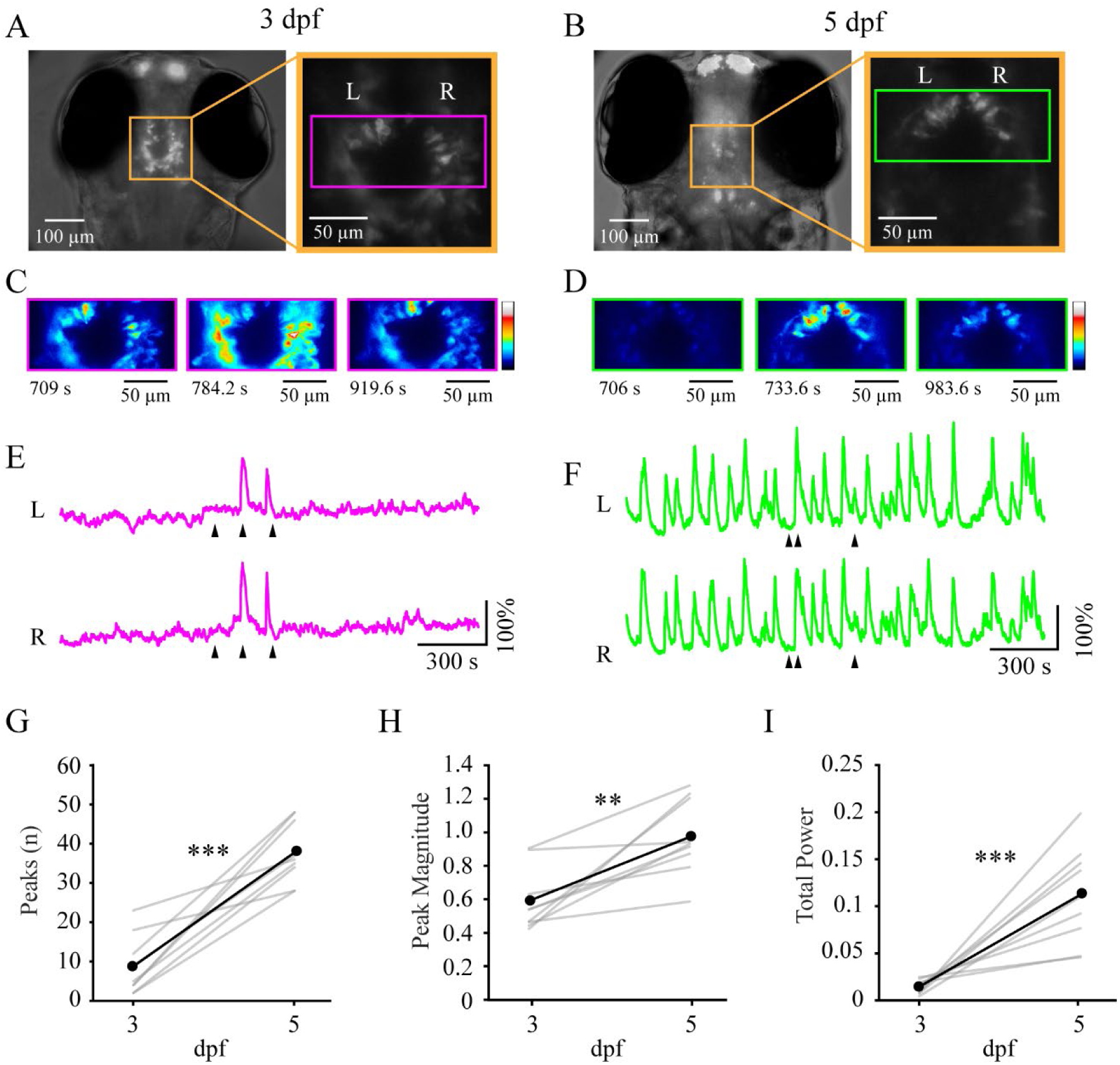
DDN/PTac neuronal activity is greater in mature larvae. (**A-B**) Representative example of an individual transgenic larva (*th:Gal4^m1233^;UAS:GCaMP6s^nk13a^*) expressing GCaMP6s in dopaminergic neurons (orange boxes) imaged at both 3 dpf (A) and 5 dpf (B). Insets: Orange boxes indicate magnified view of GCaMP6s-positive neurons in the ventral diencephalon, while magenta and green boxes indicate regions of interest for calcium imaging of identified DDN/PTac neurons shown in panels C and D. Left (L) and Right (R) cell groups are indicated above magenta box. (**C-D**) Pseudocolored GCaMP6s fluorescence panels correspond to the time points (indicated by arrows below each ΔF/F traces (E and F, respectively)). Color indicates fluorescence intensity. (**E-F**) ΔF/F traces for DDN/PTac neuron groups (L and R) recorded from an individual larva at both 3 dpf (E) and 5 dpf (F). (**G**) Plot of the mean number of calcium signal peaks detected in immature (3 dpf) and mature (5 dpf) larvae. Gray lines indicate within subjects repeated measures, while black lines represent the mean (n = 9). (**H**) Plot of the mean magnitude of calcium signal peaks in in immature (3 dpf) and mature (5 dpf) larvae. Gray lines indicate within subjects repeated measures, while black lines represent the mean (n = 9). (**I**) Plot of the mean total power of calcium signals in in immature (3 dpf) and mature (5 dpf) larvae. Gray lines indicate within subjects repeated measures, while black lines represent the mean (n = 9). Asterisks indicate significant differences at **p < 0.01 and ***p < 0.001.

**Table 4.**
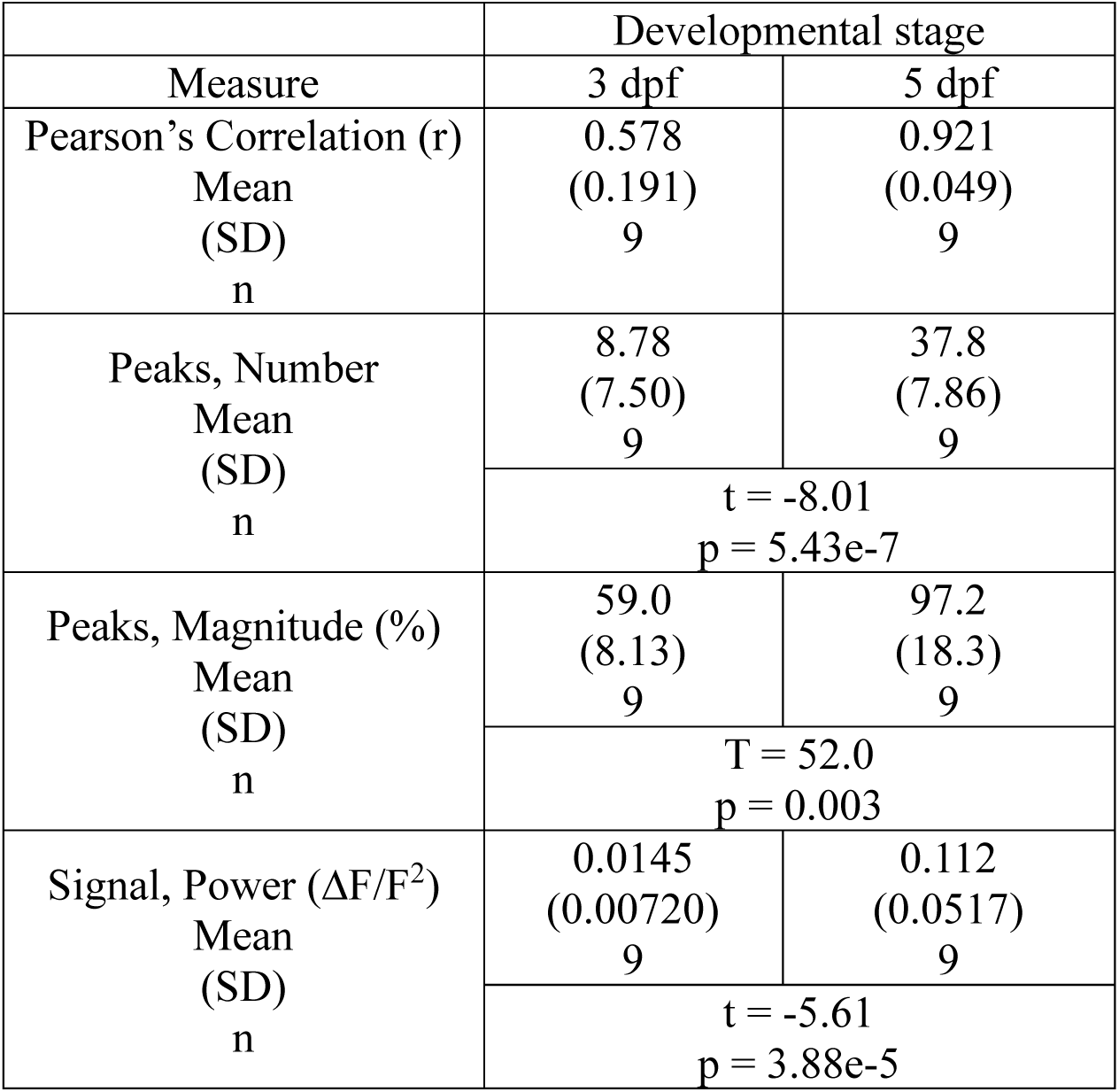
Within-subjects comparisons of calcium signal measures during spontaneous locomotor activity between immature (3 dpf) and mature (5 dpf) larvae. Values reported as mean (SD).

## Discussion

The goal of this study was to characterize the functional and molecular mechanisms responsible for the behaviorally relevant refinement of locomotor activity during development in a vertebrate model system, larval zebrafish. We hypothesized that the coarse locomotor activity produced by immature (3 dpf) larvae was due, in part, to a lack of D4Rs present in locomotor-related spinal neurons, a lack of activity in the DDNs, a lack of endogenous DA release from the DDNs in the spinal cord, or a combination of these. The work presented here demonstrates that the developmental transformation of locomotor activity correlates with an increase in gene expression of all D2-likeR subtypes, an increase in global dopamine available for release, and an increase in activity state of a group of DDNs that project to the spinal cord.

### The spinal locomotor circuit is less functionally organized in immature larvae

Although spontaneous locomotion in both free- and fictive-swimming preparations produced by immature larvae is less frequent, more erratic, and of longer duration than those produced by mature larvae (Lambert et al., 2012), the functional differences of the spinal locomotor circuit before and after the transformation of locomotor activity have not been characterized. We demonstrate here that some properties of NMDA-induced locomotor activity changed during the developmental transformation. First, we showed that the proportion of immature larvae that produced episodically organized locomotor activity was significantly lower than the proportion of mature larvae, indicating that the immature spinal locomotor circuit is functionally less well developed. Second, although a proportion of immature larvae produced locomotor activity with an EO score greater than 0.75, the scores were significantly lower compared to mature larvae. Again, this indicates that the spinal locomotor circuit is functionally less well developed. Third, the episode durations produced by immature and mature larvae were not significantly different. This result is consistent with our previous result that demonstrated NMDA-induced long duration episodes were produced in both spinalized immature and mature larvae (Lambert et al., 2012). Finally, the number of bursts generated by immature larvae were significantly greater compared to mature larvae. Overall, these results converge to reveal a functional difference in the spinal locomotor circuit between immature and mature larvae.

### D4Rs are present and functional in immature larvae

Previously, we demonstrated that dopaminergic signaling through inhibitory dopamine receptors was necessary to maintain the mature locomotor phenotype following the developmental transformation of locomotor activity (Lambert et al., 2012). Therefore, we reasoned that the coarse locomotor activity (swimming) produced by immature larvae may be due to a lack of functional D4Rs present in locomotor-related spinal neurons.

First, we confirmed that NMDA-induced episodically organized locomotor activity was sufficiently stable over time in immature larvae to measure potential changes in response to application of D2-likeR agonists. Neither the duration of episodes, the EO scores, nor the total number of bursts were affected by the duration of NDMA application. These results confirmed that NMDA-induced episodically organized locomotor activity produced by immature larvae was sufficiently stable over time (∼60 min) to assess the effects of pharmacological perturbation of dopamine receptors. Further, since DMSO was used as a vehicle for the D4R agonist, we tested whether vehicle administration had an effect on NMDA-induced locomotor activity. Application of vehicle did not produce significant differences in episode duration, episode organization, or the total number of bursts.

Next, we tested the hypothesis that inhibitory dopamine receptors were present and functional in immature larvae. Application of a broad D2-likeR agonist did not significantly affect either episode duration or number of bursts during NMDA-induced fictive swimming in immature larvae. Although episode duration was reduced in immature larvae, the effect was not significant. Likely, the statistical power was insufficient since the sample size (n = 5) was small for this group, which was reflected by the low success rate (15%; 35 out of 236 experiments) for generating NMDA-induced episodically organized locomotor activity in immature larvae. Importantly, the EO scores significantly decreased. Application of a specific D4R agonist significantly reduced both episode duration and EO score during NMDA-induced fictive swimming in immature larvae without affecting the number of bursts. Our overall interpretation of these results is that activation of D2-likeRs or D4Rs in immature larvae does not function to reduce episode duration. Rather, it functionally disrupts the organization of bursts into episodes but does not, overall, inhibit NMDA-induced locomotor activity. Unfortunately, we were unable to disambiguate D2R and D3R subtype function in immature larvae because of the high affinity of the D2-like agonist (quinpirole) to both D2Rs and D3Rs (Seeman and Van Tol, 1994).

### Gene expression of all D2-likeRs is greater in mature larvae

Our previous work reported that the developmental transformation of locomotor activity, was dependent on D2-likeR and D4R signaling (Lambert et al., 2012). In addition, we demonstrated in this study that D4Rs were both present and functional in immature larvae. Therefore, we tested the hypothesis that developmental changes in gene expression levels of all D2-likeRs in the spinal cord function to support the developmental transformation of locomotor activity. The qRT-PCR analysis revealed that all D2-like receptor subtype transcripts were present in immature larvae, and all, except *drd2l*, were significantly greater in mature larvae. The relative increase in gene expression of these receptor genes correlates with the developmental transformation of locomotor activity, which supports our hypothesis that an increase in D2-like receptor subtypes in mature larvae is, at least, one mechanism that contributes to the developmental transformation of locomotor activity. Although we did not directly measure protein levels for D2-like receptor subtypes, our functional data indicate that these receptors are present and functional in immature larvae.

Characterization of the spatial distribution of D2-like receptor subtypes on identified locomotor-related spinal neurons would provide insights into the functional mechanisms responsible for the developmental transformation of locomotor activity. Previous studies identified candidate locomotor-related neurons that expressed D2-likeRs. One study demonstrated that V3 spinal interneurons were hyperpolarized in response to D2-likeR agonist (quinpirole) application in mice (Sharples et al., 2020). A second study showed that DDN projections (DDT) likely synapse on spinal motor neurons and that these synapses co-localized with D4Rs in larval zebrafish (Son et al., 2020). Therefore, in future experiments, we plan to identify locomotor-related spinal neurons that express D2-like receptor subtypes in both immature and mature larvae by mapping their locations and relative abundance using third-generation *in situ* hybridization chain reaction.

### More dopamine is present in mature larvae

We have demonstrated that D4R subtype is present and functional in the spinal locomotor circuits of immature larvae, and that gene expression of all D2-like receptor subtypes are increased in the bodies of larval zebrafish across the developmental transformation of locomotor activity. Next, we tested the hypothesis that an increase in DA levels correlated with the transformation of locomotor activity. Whole-body (global) DA levels increased significantly during the developmental transformation. Although our data do not directly demonstrate that more DA is present in nervous tissue of mature larvae, it provides evidence that more DA is present globally in mature larvae.

### DDNs are more active in mature larvae

We reasoned that since a greater amount of dopamine was present in mature larvae, then an increase in activity in DDN/PTac neurons in mature larvae would increase the yield of dopamine release on spinal locomotor circuits. Thus, we used DDN/PTac activity as a proxy for dopamine release to test the hypothesis that increased DDN/PTac activity from the immature to the mature stage of larval development correlated with the locomotor transformation.

Although the DDN/PTac neurons generated a range of activities in both immature and mature larvae, the measured calcium transients increased within all preparations during development. Specifically, the number of calcium transient peaks, the magnitude of calcium transients, and power measures of the calcium transients were significantly greater in mature larvae compared to immature larvae. These results support the hypothesis that increased dopaminergic signaling through DDN/PTac activity correlates with the transformation of locomotor activity during development.

DDN/PTac neurons are located in the ventral diencephalon, which necessitated calcium imaging experiments to be performed in intact larval preparations. Since DDN activity is correlated with locomotion in larvae between 4-6 dpf (Jay et al., 2015; Reinig et al., 2017), the difference in DDN/PTac neural activity between larvae at 3 dpf (immature) and 5 dpf (mature) could be a consequence of increased locomotor activity driven by higher-order centers in mature larvae. Regardless, greater DDN/PTac activity in mature larvae likely corresponds to greater release of DA on spinal locomotor circuits since greater levels of DA are present.

In conclusion, this work provides convergent evidence for multiple mechanisms underlying the developmental transformation of locomotor activity in vertebrates. We showed that the spinal locomotor circuitry is more functionally organized after the developmental transformation, and that D4Rs were both present and functional in the spinal locomotor circuits of, at least, a subset of immature larvae. Importantly, gene expression of all D2-like receptor subtypes, levels of dopamine, and activity of diencephalic dopaminergic neurons were significantly greater in mature larvae compared to immature larvae. Overall, the integration of these results indicates an important developmental role for dopaminergic signaling in the refinement of locomotor activity in vertebrates.

